# Local Polyploidy Phenomenon in *Escherichia coli* and its significance in genetic engineering

**DOI:** 10.1101/335448

**Authors:** Fayu Yang, Liwei Guo, Kenao Lyu, Jiao Zhang, Yunpeng Yang, Huiyan Wang, Huan Yu, Dele Guo, Tingting Ding, Chaoyong Huang, Huiyong Ren, Xiaoyan Ma, Yu Yang, Yi-xin Huo

## Abstract

Classic Helmstetter & Cooper model asserted that the multifork phenomenon in the process of replication. However, the impacts of the multifork on the evolution and genetic engineering are still vague. Here, we employed CRISPR/Cas9 technology to knock-out eighteen *Escherichia coli* chromosomal fragments (over 50 kb) that represent all areas of the chromosome. We demonstrated that a single cell could have wild-type, color-reporter, and antibiotic-resistant genes simultaneously in the same locus of the different branches of the duplication forks after multiple rounds of deletions and replacements. This phenomenon that a single cell had different genotypes in its local polyploid chromosomes, which was similar to eukaryotic heterozygote, was named as local polyploidy. Under a defined selective pressure condition, offspring cells containing at least a copy of conditionally beneficial mutation could be enriched, and other alleles could be kept silently and peacefully in the duplication fork(s) of the same cell. The significance of this phenomenon in the genetic engineering was discussed.

## 1. Introduction

Bacterial chromosome replication is best investigated in model bacteria, *Escherichia coli*(1, 2) and *Bacillus subtilis*(3, 4). A circular bacterial chromosome generates two replication forks, replicating chromosome bidirectionally from the “Origin” until the “Terminus”. Each half of the chromosome replicated by one of the two replication forks is called a “Replichore”(5-7). However, some cells initiated new rounds of chromosome replication before completing the previous one, resulting in two, four or even eight rounds of replication proceeding simultaneously. This phenomenon, which was termed “multifork replication”(8), was formalized by Cooper and Helmstetter in 1968(9). Notably, Cooper and Helmstetter’s model illuminated that cells could balance the largely constant rates of replication fork progression with the nutrient-dependent changes in mass doubling time, by initiating replication and dividing more frequently when growing faster(9, 10) (**Supplementary Figure 1**). Although the development of bacterial quantitative physiology and fluorescence microscopy can infer the presence of multifork, the size and the life cycle of each replication fork in the multifork remains vague(11-15).

Multiple copies of chromosomal fragments will exist in local areas if the phenomenon of bacterial multifork exists. If deletion or replacement of targeted chromosomal fragment did not occur in all copies of the chromosome in the replication fork, a local polyploidy phenomenon will occur, leading to the behavior segregation among offspring cells. It is interesting to know whether a single bacterial cell containing different alleles in the same locus could mimic the fate of an eukaryotic heterozygous cell. What are the impacts of the multifork triggered local polyploidy or “heterozygote” on the evolution and genetic engineering?

The emergence of CRISPR/Cas9 technology provides us a possible tool to investigate the above puzzles. In the CRISPR/Cas9 system, the Cas9 nuclease directly introduces double strand breaks (DSBs) at a position 3 bp upstream of the DNA double strand containing PAM (Protospacer adjacent motif) with the guidance of two noncoding RNAs: CRISPR RNA (crRNA) and trans-activating crRNA (tracrRNA)(16-20). Since 2013, CRISPR/Cas9 system has been successfully applied to multiple species, including bacteria(21-24), yeast(25, 26) and mammalian cells(27, 28). It has been widely used in *E. coli* for generating site-directed mutations, codon substitution, fragment deletion, and other precise gene modifications (29-31). It is well known that homologous recombination (HR) is a major DNA repair pathway in prokaryotic organisms and is particularly important in such bacteria lacking non-homologous end joining (NHEJ) such as *E. coli*(32). In *E. coli*, the repair of DSBs needs an exogenous donor DNA through HR(33, 34). However, due to the long spatial distance between the two incision ends of a large truncation, its HR based repair by a small donor DNA remains challenging and could be very inefficient without an overexpressed recombination system.

In this study, CRISPR/Cas9 technology was utilized to challenge the replacement of chromosome fragments over 50 kb in *E. coli* without overexpressing any endogenous or exogenous recombination system. Due to the low efficiency, it is possible that only part of the chromosome copies in the branches of a multifork could be deleted and replaced in one bacterial generation (**Fig. 1**). Reporter gene was used to replace the deleted chromosome fragments at different locations to construct a reporter system. Here, we demonstrated that the same locus at the different copies of chromosomes in the replication fork of a single cell could be edited into different genotypes by knocking-in color reporter genes, antibiotic resistance genes, etc. The large fragment deletion experiments were performed at eighteen different locations on the chromosome, representing all areas of the chromosome. Local polyploidy or “heterozygote” phenomenon were observed in all engineered strains, indicating that the size and life span of bacterial replication fork may be greater than our traditional belief, and that multifork may exist not only during the rapid growth period.

**Figure 1:**
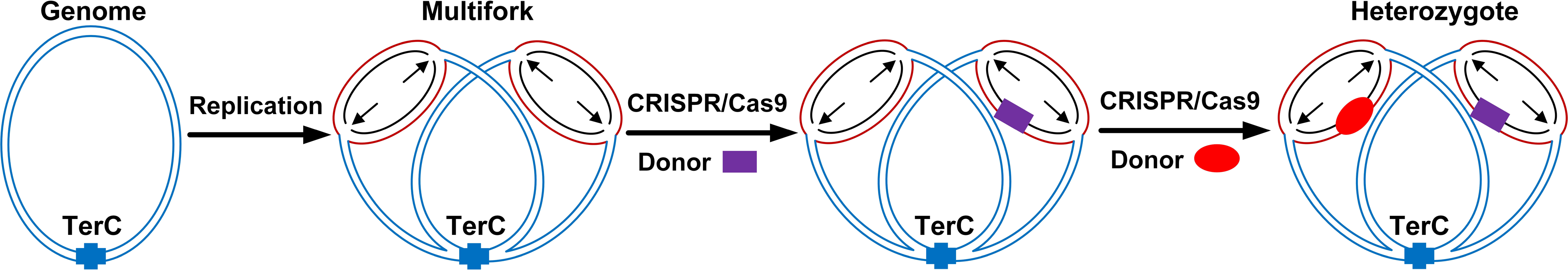
The hypothesis of multifork to local polyploidy or “heterozygote” during the bacterial reproduction. Multifork replication refers to the re-initiation of a new round of replication per chromosome before the first round of termination, ensures that at least one round of replication is finished before cytokinesis, to guarantee that each daughter cell receives at least one complete genome. The same locus at the different copies of chromosomes in the replication fork of a single cell could have different genotypes, such as wild-type, PPRG, and KRG, forming a local polyploidy or “heterozygote” phenomenon. These three genotypes could be segregated to different progeny cells in different combinations, so that the offspring cells have different phenotypes and exhibit different growth conditions under different environmental conditions.

The mixed genotypes could be segregated into offspring cells in different combinations, so that those offspring cells could have different phenotypes (**Fig. 1**) and exhibit different growth behaviors under different environmental conditions. Genetic engineering that is temporarily non-beneficial or even harmful to the cells under non-stressed conditions could provide cells the resistance toward an occasional environmental stress. The engineered cells could be enriched to become dominant genotype under the corresponding stress. When the corresponding stress disappeared, the engineered strains lost their competitive advantage and the population harboring that mutant decreased due to its fitness cost. These results shed new light on a novel possibility of strain degeneration in the field of genetic engineering.

## 2. Materials and Methods

### 2.1 Strains and Culture Conditions

The *E. coli* K-12 strain Top10 (*BM* Top10, Biomed) was used as the host strain for molecular cloning. The genotype of the strain Top10 is F-mcrAΔ (mrr-hsdRMS-mcrBC) Φ 80lacZ Δ M15 Δ lacX74recA1ara Δ 139 Δ (ara-leu) 7697galUgalKrpsL (Starr) endA1nupG. The mutant recA1 protein has a greatly reduced single-stranded DNA-dependent ATPase activity at pH 7.5 compared to the wild-type protein (35). *E. coli* strains Top10 was taken as the host for genetic engineering using CRISPR/Cas9. Strains for cloning were grown in Luria-Bertani (LB) medium (10 g/l tryptone, 5 g/l yeast extract, and 10 g/l NaCl) supplemented with appropriate antibiotic (ampicillin (100 μg/ml), kanamycin (25 μg/ml) and chloramphenicol (50 μg/ml)).

### 2.2 Plasmid construction

The plasmid pCas9cur encoding Cas9 protein was constructed originally by *Li, Y. F. et al*(29). The Flp gene was introduced into the plasmid pGRB(29) to obtain the plasmid pFlp-sgRNA. To generate pJET-sgRNA-pCI, the sgRNA-pCI fragment was synthesized by PCR-based accurate synthesis (PAS) method. Plasmid pJET-sgRNA-pCI was constructed by ligating the sgRNA-pCI fragment with the vector pJET1.2 (CloneJET PCR Cloning Kit, Thermo Fisher Scientific). Plasmid DNA was isolated by standard techniques. All DNA sequences in the constructs were confirmed by sequencing. All primers used in plasmids construction could be found in **Supplementary Table 5**.

To construct sgRNA plasmid, a set of primers were used to PCR amplify the pFlp-sgRNA backbone. The 20 bp spacer sequence designed for each target was synthesized in primers. The PCR product was then self-assembled using Gibson Assembly to obtain the desired gRNA plasmid **(Supplementary Figure 3 and Table 2)**.

Donor dsDNA usually contained 300∼500 bp homologous arm on each side (5’ homologous arm had 500 bp and 3’ homologous arm had 300 bp). To construct donor dsDNA, two homologous arms and the sequence (the prancer purple reporter gene (PPRG) or kanamycin resistance gene (KRG)) to be inserted were separately amplified and were then fused together by OE-PCR. The PPRG come from DNA2.0 (CPB-37-441). We measured the growth rates of two Top10 derivatives with or without the PPRG overexpression (**Supplementary Figure 34**). The results showed that the growth rates of the two Top10 derivatives were similar, in agreeing with the conclusion that the PPRG did not affect the growth of the host cells (**(https://www.atum.bio/eCommerce/catalog/datasheet/529).**PCR products were cloned into the vector pJET1.2 (CloneJET PCR Cloning Kit, Thermo Fisher Scientific). The plasmids were digested by appropriate restriction enzymes, usually by NotI and PstI unless otherwise noted. Gel purification of the enzyme-digested PCR products prior to electroporation is necessary. All primers used in donor dsDNA construction could be found in **Supplementary Table 3**.

### 2.3 Oligonucleotides and PCR

All primers were designed by Integrated DNA Technologies (IDT) (http://sg.idtdna.com/sessionTimeout.aspx). Oligonucleotide sequences were listed in **Supplementary Table 2**,**Table 3, Table 4 and Table 5**. PCR was performed with 1 μl of 2.5 U/μl TransStart FastPfu Fly DNA Polymerase (TRANCEGEN) in 50 μl with 1×;FastPfu Fly Buffer, 0.2 mM dNTP mix (TRANCEGEN), 0.2 μM of each primer and a program of: 95 °C, 3 min; 32 cycles of (95 °C, 20 s; 58 °C, 15 s; 72 °C, 1 min) unless otherwise noted.

### 2.4 Genome editing procedure in *E. coli*

Top10pCas9cur competent cells were generated as previously described(36). 200 ng donor dsDNA and 100 ng sgRNA plasmid were added in each reaction. After transformation, 500 μl of LB were immediately added into the cells. After 1 h recovery, the cells were centrifuged to remove the supernatant. 1 ml LB (Cm + Amp) was added into the tube. Cells were recovered for 3 h prior before 200 μl of the cells were plated. Plates were cultivated at 37 °C for 48 h.

### 2.5 The dilution-plating method

This method was typically used to separate microorganisms contained within a small sample volume (eg. a single colony), which was spread over the surface of an agar plate, resulting in the formation of discrete colonies distributed evenly across the agar surface when the appropriate concentration of cells was plated(37). A single colony was picked and diluted in 1000 μl ddH_2_O or LB broth. After mixed gently, 50 μl of diluted cells were plated on the plates containing different antibiotics. The plates were cultivated at 37 °C. The numbers of colonies formed were manually counted.

### 2.6 Isobutanol tolerance assay

Isobutanol tolerance was determined by calculating the numbers and ratios of the white or purple cells, and by measuring the OD_600_. For OD_600_ measurement, 1% (vol/vol) of the overnight culture was inoculated in 200 ml LB medium in 500-ml baffled shake flasks and grew at 37 °C until early exponential phase (OD_600_, 0.45–0.5). 30 ml of culture was then inoculated into a 250-ml baffled shake flask, followed by the addition of isobutanol with desired concentrations. We used parafilm to wrap the cap of the test tube to minimize the evaporation. The growth of cells was sampled and monitored by OD_600_ measurement. The ratio of OD_600_ at 48 and 0 h was used to determine the tolerance. Viable cell counting was performed after OD_600_ measurements. After dilution by LB broth, cells were plated on an LB (no antibiotic) plate and an LB (Cm + Amp) plate. Plates were incubated at 37 °C for 48 h. The numbers of colonies developed on the plates were counted.

### 2.7 Purification of Genomic DNA and Sequencing

Genomic DNA was purified from cells using the bacterial genomic DNA purification spin columns (TIANGEN). The flanking sequences of the reporter genes were PCR amplified. The PCR products were purified and then sequenced by Synbio Tech (sequencing primers were shown in **Supplementary Table 4**).

## 3. Results and Analysis

### 3.1 The presence of bacterial multifork phenomenon demonstrated at the single-cell level

CRISPR/Cas9 dependent large fragments deletion and replacement in *E. coli*, a species without NHEJ system, relied on the repair of DSB by HR(21, 32, 38). There was no previous report for HR-dependent deletion and replacement for any over 50 kb fragment in *E. coli*. Here, we challenged to delete over 50 kb fragments in an *E. coli* strain Top10, which had even greatly reduced recombinase activities(35) (see also materials and methods). Since the efficiency of HR was low in Top10, we assumed that it was possible that only part of the chromosome copies in the branches of a multifork could be deleted and replaced in one bacterial generation. To testify this hypothesis, we created a reporting system by knocking-in the Prancer purple reporter gene (PPRG) (See **Fig. 2a**) at eighteen different locations of the *E. coli* chromosome, spanning all the areas of the chromosome (four knock-in locations next to the OriC and four knock-in locations next to the Terminus) (**Supplementary Figure 2 and Table 1**). For example, a pair of sgRNAs were designed on both sides of the gene sequence for the chromosomal fragment (56 kb) within the range of 1,588,791-1,645,053 of the *E. coli* genome. An expression vector containing the above sgRNAs was constructed and named as pFlp-sgRNA_56 kb, carrying the ampicillin (Amp) resistance marker. PPRG with homologous arm was designed according to the target gene sequence (**Fig. 2a**). The pCas9cur plasmid with the Cas9 protein encoding gene and the chloramphenicol (Cm) resistant marker was introduced into the Top10 strain, generating the Top10pCas9cur strain. The linearized donor DNA (Donor-PPRG) containing the PPRG, and the pFlp-sgRNA_56 kb or its negative control were co-transformed into Top10pCas9cur competent cells (**Fig. 2b, Supplementary Figure 3, Table 2 and Table 3**). Only transformants containing both of the plasmids pCas9cur and pFlp-sgRNA_56 kb could resist both of the Cm and Amp antibiotics. If Cas9 protein did not work, the cells will remain white. If Cas9 protein worked well, DSB could be introduced into the *E.coli* chromosome. If the DSB could not be repaired, the cells will die. If the DSB could be repaired by Donor-PPRG through HR, the cells will turn purple owing to the purple protein expressions.

**Figure 2:**
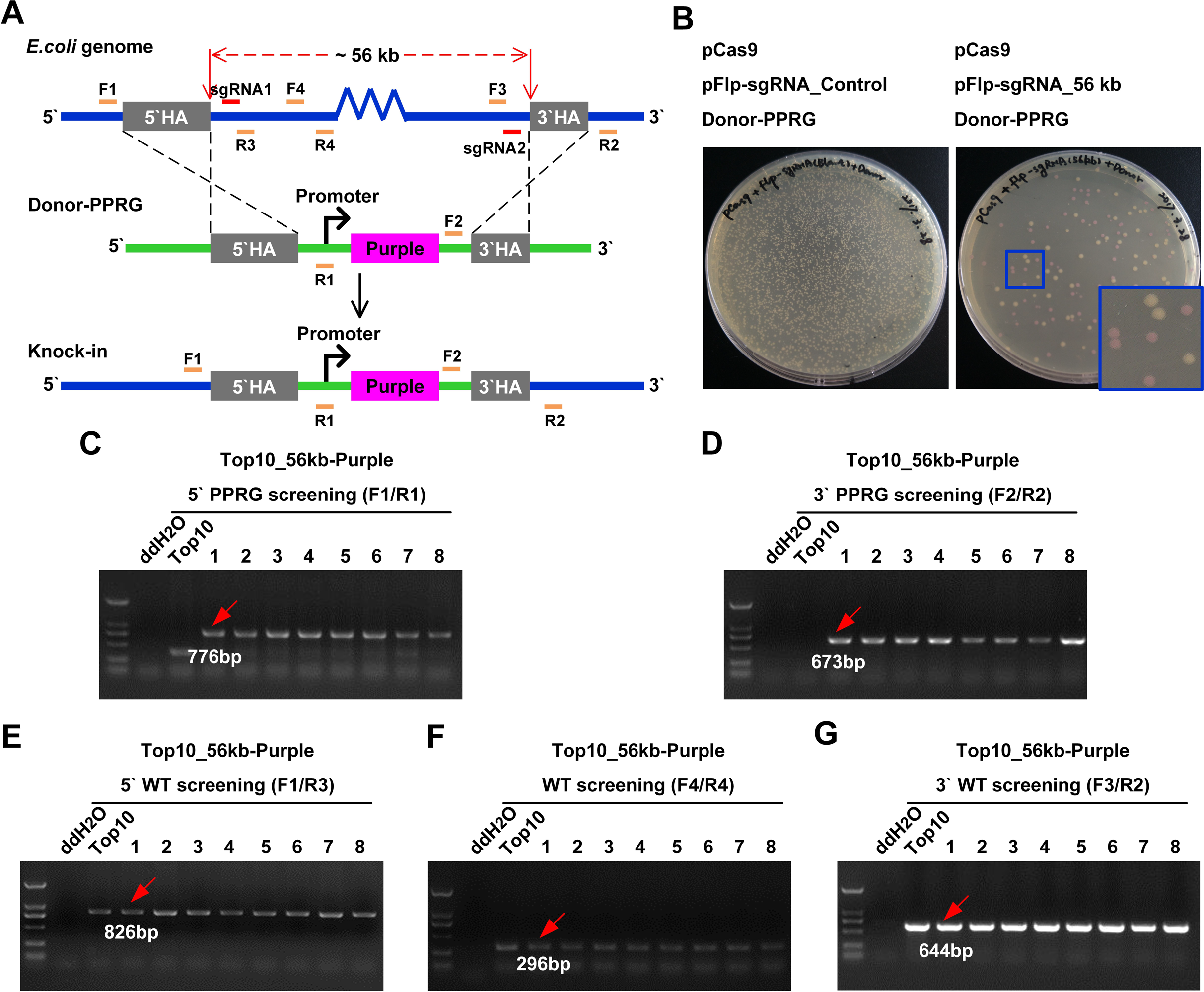
Generation of 56 kb <u>(NO. 9)</u> fragment deletion and the Prancer purple reporter gene (PPRG) knock-in at the genome in *E. coli*. A. Schematics of the donor DNA and targeting strategy for CRISPR/Cas9-mediated homology recombination, which integrated the PPRG into the chromosome at the position of the 56 kb fragment deletion. Black lines indicated sections of homology between the genomic locus and the donor DNA. Positions of PCR primers used for detecting of the PPRG knock-in were shown. B. Top10pCas9curcompetentcellswereco-transformedwith pFlp-sgRNA_Control/56 kb (100 ng) and Donor-PPRG DNA (200 ng), and images were obtained at 48 h post transformation (PPRG knock-in colored in purple; WT colored in white). C. Recombination screening of left arm (5’ homologous arm) by PCR. Genotyping of the 56 kb fragment deletion in the genome and the screening of PPRG knock-in candidate colonies with F4/R4 primers (at least on PCR products with correct size were purified and verified by sequencing, results not shown). See also **Supplementary Table 4.** D. Recombination screening of left arm (3’ homologous arm) by PCR. Genotyping of the 56 kb fragment deletion in the genome and the screening of PPRG knock-in candidate colonies with PCR (F5/R5 primers) (at least on PCR products with correct size were purified and verified by sequencing, results not shown). See also **Supplementary Table 4.** E. Wild-type screening of the 5’ boundary sequence of wild-type 56 kb fragment by PCR (F1/R3 primers) (at least on PCR products with correct size were purified and verified by sequencing, results not shown). See also **Supplementary Table 4.** F. Wild-type screening of the inside sequence of the wild-type 56 kb fragment by PCR (F4/R4 primers) (at least on PCR products with correct size were purified and verified by sequencing, results not shown). See also **Supplementary Table 4.** G. Wild-type screening of the 3’ boundary sequence of the wild-type 95 kb fragment by PCR (F3/R1 primers) (at least on PCR products with correct size were purified and verified by sequencing, results not shown). See also **Supplementary Table 4.**

When plated with a diluted culture, a single colony was developed from a single transformant. This method was typically used to separate microorganisms contained within a small sample volume (eg. a single colony), which was spread over the surface of an agar plate, resulting in the formation of discrete colonies distributed evenly across the agar surface when an appropriate concentration of cells was plated (37). Single colonies appeared on LB (Cm + Amp) plates after 12 to 24 h incubation at 37 °C. Some colonies turned purple after an additional day (**Fig. 2b**), named as Top10_56 kb-Purple. Eight purple single colonies were randomly picked and verified by PCR targeting the 5’ and 3’ ends of the PPRG, respectively, to confirm that the 56 kb fragment of the chromosome was replaced by the PPRG. PCR results showed that all eight colonies were positive colonies (**Fig. 2c, 2d and Supplementary Table 4**) and the sequencing results also confirmed that the PPRG did exist at the target position. We designed primers outside and inside of the knockout construct to check the boundaries (**Fig. 2a**). Further PCR and sequencing results confirmed that all of the purple single colonies also contained the wild-type chromosomal fragment at the same target location where the PPRG was knocked-in (**Fig. 2e, 2f, 2g and Supplementary Table 4**). To further verify the above phenomenon, we constructed gene replacements for large fragments over 50 kb at other seventeen locations on the chromosome. As expected, the PCR verification results were similar to that of the strain with the 56 kb fragment deletion (**Supplementary Figures. 4-20**). Negative and positive controls were included in all of ninety PCR gel pictures. All results confirmed that the WT region and PPRG insert co-existed in every purple single colony we tested, no matter of the location of the large fragment deletion and the following PPRG knock-in. Our experiment design and sequencing results ruled out the possibility of a tandem positioning of the wild-type and the PPRG fragment **(Fig. 2 and Supplementary Figures. 4-20)**. Both wild-type and PPRG genotypes were kept intact during the process in which a single colony was developed from an engineered cell. The observations strongly indicated that only part of the chromosomal alleles in the replication fork of each engineered cell were replaced by the PPRG while the other alleles maintained their wild-type genotype.

### 3.2 The presence of bacterial local polyploidy or “heterozygote” phenomenon demonstrated at the single-cell level

The above results strongly indicated that both WT and PPRG co-existed in the same transformant. If the local polyploid or “heterozygous” cells did exist in *E. coli*, and the CRISPR/Cas9 system could continue editing the copy of the wild-type fragment in the replication fork. Thus, we would obtain the strains containing both purple and antibiotic resistance phenotypes by adding an antibiotic resistance marker into the donor DNA. If successfully conducted, a single cell could contain three different genotypes of wild-type, PPRG and an antibiotic resistance marker at the same locus of the different copies of chromosomes inside the replication fork, and become a local polyploidy or “heterozygous” cell. To verify this hypothesis, we used the kanamycin resistance gene (KRG) as part of a donor DNA to replace the 56 kb fragment at the same locus in the different chromosome copies of the genome of purple bacteria Top10_56 kb-Purple as described above. The genome copy containing the PPRG could not be further modified since the sgRNA binding site has been deleted when digested by Cas9 protein during the gene editing. As shown in **Fig. 3a**, we obtained strains containing three different genotypes of wild-type, PPRG, and KRG in the same locus of different chromosome copies in the replication fork. Here, Donor KRG, a linearized repair template containing KRG and homologous arm, was constructed according to the target gene sequence. The repair template was introduced into the Top10_56 kb-Purple competent cells of the engineered purple strain. The KRG was inserted into the genomic DNA by HR (**Fig. 3b, Supplementary Figure 21 and Table 3**), which was constructed to replace the multiple copies of the 56 kb genomic fragments in the replication fork that could express both the PPRG and KRG. The integration of KRG in the bacterial chromosome provided the cells with the resistance to the antibiotic kanamycin (Km). After 48 h incubation at 37 °C on the LB (Cm + Amp + Km) plates, colonies were obtained which should contain both PPRG and KRG (**Fig. 3c**). Eight colonies with light purple color were selected and verified by PCR at the 5’ and 3’ ends of the KRG, respectively, to confirm that the large fragment of the chromosome fragment was replaced by the KRG. PCR results showed that all 8 colonies were positive colonies (**Fig. 3d and Supplementary Table 4**) and the sequencing results also confirmed that the KRG was knocked into the targeted position (**Supplementary Figure 22 and Table 5**). The strains were named as Top10_56 kb-Purple/Km. After that, the PPRG was verified by PCR at the 5’ and 3’ ends of the PPRG, respectively, to confirm that the PPRG was still present in the positive colonies. PCR results showed that all eight colonies contained the PPRG (**Fig. 3e and Supplementary Table 4**).

**Figure 3:**
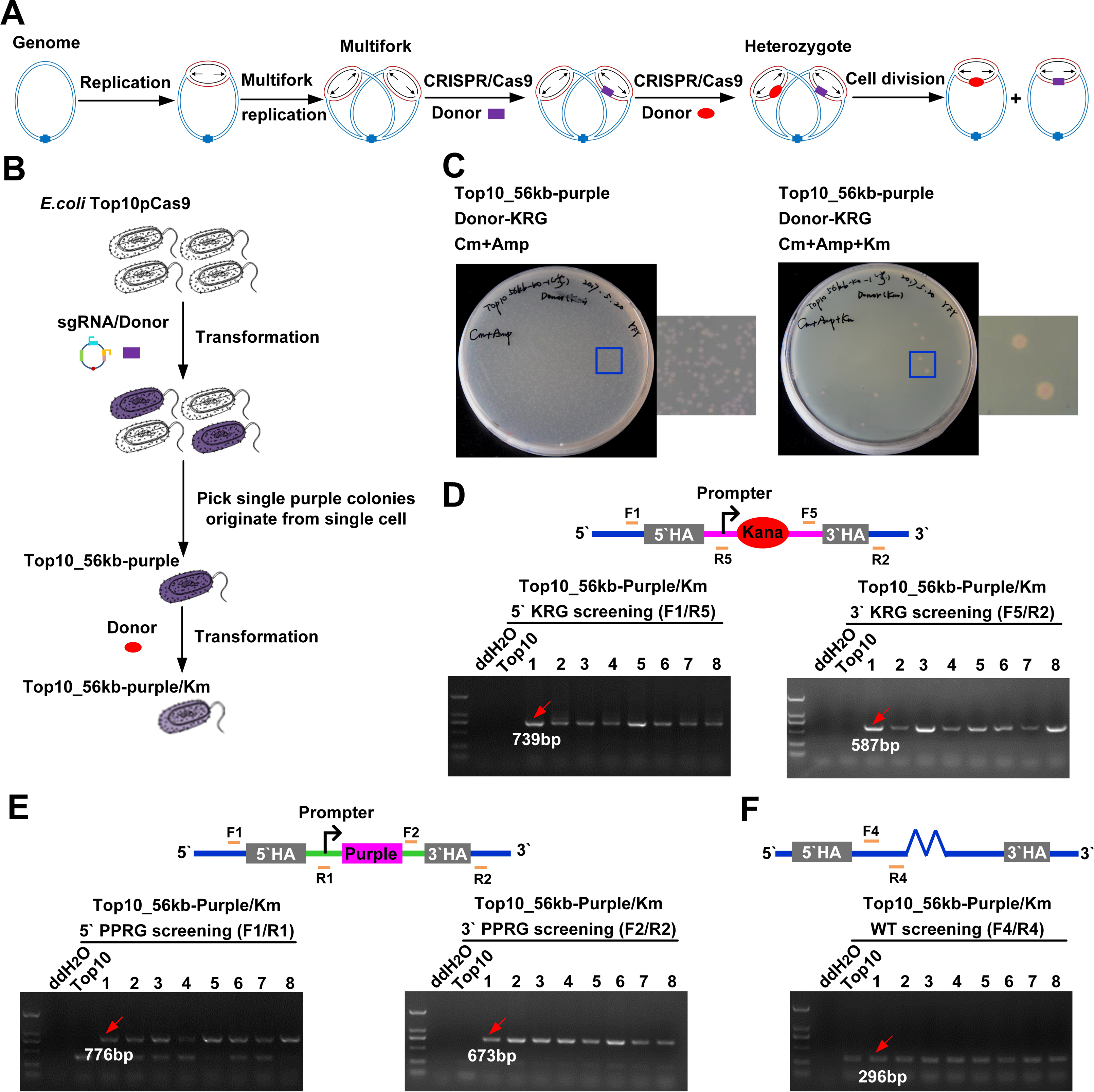
Multifork to bacterial local polyploidy or “heterozygote” triggered by iterative genome editing. a. Schematic of multifork to bacterial local polyploidy or “heterozygote” triggered by iterative genome editing. A single cell could contain three different genotypes of wild-type, PPRG, and KRG at the same locus of the different copies of chromosomes inside the replication fork, and became a local polyploidy or “heterozygote”. After cells divided, these genotypes could be allocated into the daughter cells in different combinations, so that the daughter cells have different phenotypes. b. Step-by-step schematics of the iterative genome editing. c. Top10_56 kb-Purple competent cells were transformed with Donor-KRG DNA (200 ng), and images were obtained at 48h post transformation. See also **Supplementary Figure 12.** d. KRG knock-in screening of left arm (5’ and 3’ homologous arm) by PCR. Genotyping of the 56 kb fragment deletion in the genome and the screening of PPRG knock-in candidate colonies with F1/R5 or F5/R2 primers (at least on PCR products with correct size were purified and verified by sequencing, results not shown). See also **Supplementary Table 4.** e. PPRG knock-in screening of left arm (5’ and 3’ homologous arm) by PCR. Genotyping of the 56 kb fragment deletion in the genome and the screening of PPRG knock-in candidate colonies with F1/R1 or F2/R2 primers (at least on PCR products with correct size were purified and verified by sequencing, results not shown). See also **Supplementary Table 4.** f. Wild-type screening of the inside sequence of the wild-type 56 kb fragment by PCR (F4/R4 primers) (at least on PCR products with correct size were purified and verified by sequencing, results not shown). See also **Supplementary Table 4.**

In order to verify whether local polyploidy or “heterozygote” colonies still contain wild-type sequence, we used primers outside the knockout construct and inside to check the boundaries on the 56 kb deletion fragment. The PCR assays showed that all eight positive colonies still contained wild-type fragment (**Fig. 3f and Supplementary Figure 22 and Table 4**). Moreover, we designed multi pairs of primers outside and inside the knockout construct to check the boundaries and the all length 56 kb fragment by PCR in *E. coli* Top10 or Top10_56 kb-Purple or Top10_56 kb-Purple/Km cells (**Supplementary Figure 23**). The results confirmed that in the WT copies the chromosome loci was still intact and was not edited.

### 3.3 Bacterial local polyploidy or “heterozygote” demonstrated by the dilution-plating assays

Every Top10_56 kb-Purple/Km cell grew on LB (Cm + Amp + Km) plates was Km resistant and should have at least one copy of the KRG in the chromosome. To confirm that three genotypes co-existed in the engineered cells, three positive single colonies (named as G2 single colony, the abbreviation of the Generation 2 single colony) of Top10_56 kb-Purple/Km from LB (Cm + Amp + Km) plates was picked.

Every G2 was developed from a transformant (named as G1 transformant). Each G2 were diluted in 1000 μl ddH_2_O, and 50 μl of cells were plated on LB (no antibiotic), LB (Cm + Amp), LB (Km) and LB (Cm + Amp + Km) plates. As explained earlier, the chromosome could not be further edited if any of the plasmid was lost due to the lack of its corresponding antibiotic pressure during the new plate incubation time. The growth of colonies was observed after incubation at 37 °C for 48 h (**Fig. 4a**). On LB (no antibiotic) or LB (Cm + Amp) plates, cells without KRG genotype could survive. Among the colonies (named as P3 single colony), about 20% were purple and the others were white (**Fig. 4b and Supplementary Figure 24b**). On LB (Km) or LB (Cm + Amp + Km) plates, only cells with KRG genotype could survive. Among the colonies (also named as G3 single colony), about 10% were light purple and the others were white (**Supplementary Figure 24b**). Every G3 single colony was developed from a single cell in a G2 single colony, while that G2 single colony was developed from a G1 transformant. Since 10-20% of G3 single colonies contained the PPRG, it was obvious that the G1 transformant contained PPRG. Since every G2 single colony was grew on LB (Cm + Amp + Km) plates, the G1 transformant and every cell in a G2 single colony should have at least one chromosome copy of the KRG. Taken together, the original G1 transformant contained PPRG and KRG simultaneously.

Furthermore, the assays in **Supplementary Figure 24b** were modified by culturing the G2 single colonies in liquid LB (no antibiotic) or LB (Km) medium for 48 h at 37 °C before plating (Details summarized in **Supplementary Figure 25a**). Results in **Supplementary Figure 25b and 25c** demonstrated that a G2 single colony originally derived from a single G1 transformant (Top10_56 kb-Purple/Km) showed phenotypic segregation after culturing under different environmental conditions. Taken together, the genome of a single cell could contain wild-type, PPRG, and KRG genotypes in the same locus of the different copies of chromosomes in the replication fork *in E. coli* (**Fig. 4c**).

**Figure 4:**
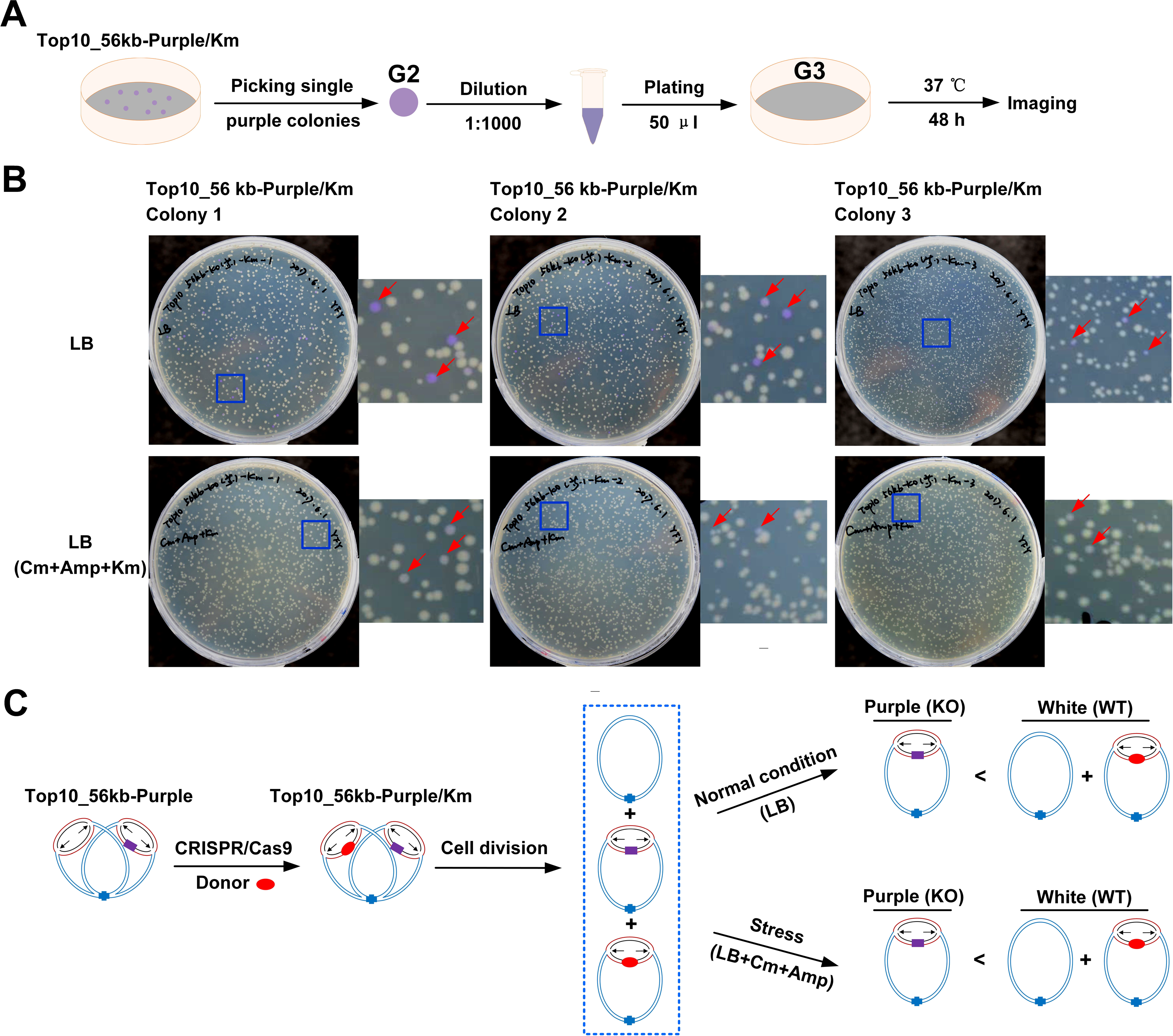
The phenomenon of bacterial local polyploidy or “heterozygote”. A An illustration of the Top10_56kb-Purple/Kana spotted on different resistance plates to test the local polyploidy or “heterozygote” phenomenon in bacteria. A single colony from LB (Cm + Amp + Km) plate was picked and diluted in 1000 μl ddH_2_O, and 50 μl of cells were plated on plates with different antibiotics and cultivated at 37 °C. B Top10_56 kb-Purple/Kana single colony could have different phenotypes in the daughter cells. Three positive colonies from LB (Cm + Amp + Km) plate was randomly picked and plated on plates with different antibiotics. After incubation at 37 °C for 48 h, it was evident that about 80% of the colonies grown on LB (no antibiotic) plates were white colonies and 20% of the colonies on the same plate were purple. Among the colonies on the LB (Cm + Amp + Km) plates, 10% of the colonies were purple colonies and 90% of the colonies were white colonies. See also **Supplementary Figure 24.** C Hypothesis of bacterial local polyploidy or “heterozygote” caused by multifork replication. A single cell genome replication forks in the same locus in the different chromosome copy can type different genes at the same time, such as wild-type, PPRG and KRG, and become a local polyploidy or “heterozygote”. Thus, in the single colonies formed by cell division and progeny single cells, both white and purple single colonies appear simultaneously (WT indicates wild-type strain; KO indicates engineered strain).

### 3.4 Advantages of local polyploidy or “heterozygote” phenomenon in bacteria in environmental stress response

In order to explore the role of local polyploidy or “heterozygote” in the bacteria evolution, we hypothesized that we could adjust the ratio of purple (engineered strain) to white (wild-type strain) colonies in the bacterial population by changing the environmental conditions. We further tested whether the shift of the environment could also lead to the replacement of dominant strains (**Fig. 5a**). By measuring the growth of Top10_56 kb-Purple under a series of growth conditions (e.g., different pH, salt concentration, temperature, isobutanol, etc.), we found that Top10_56 kb-Purple grew slower than wild type in the absence of stress, but was more tolerant to high isobutanol concentration. Therefore, the isobutanol pressure could be used as a selection pressure to enrich the engineered 56 kb deletion strains.

**Figure 5:**
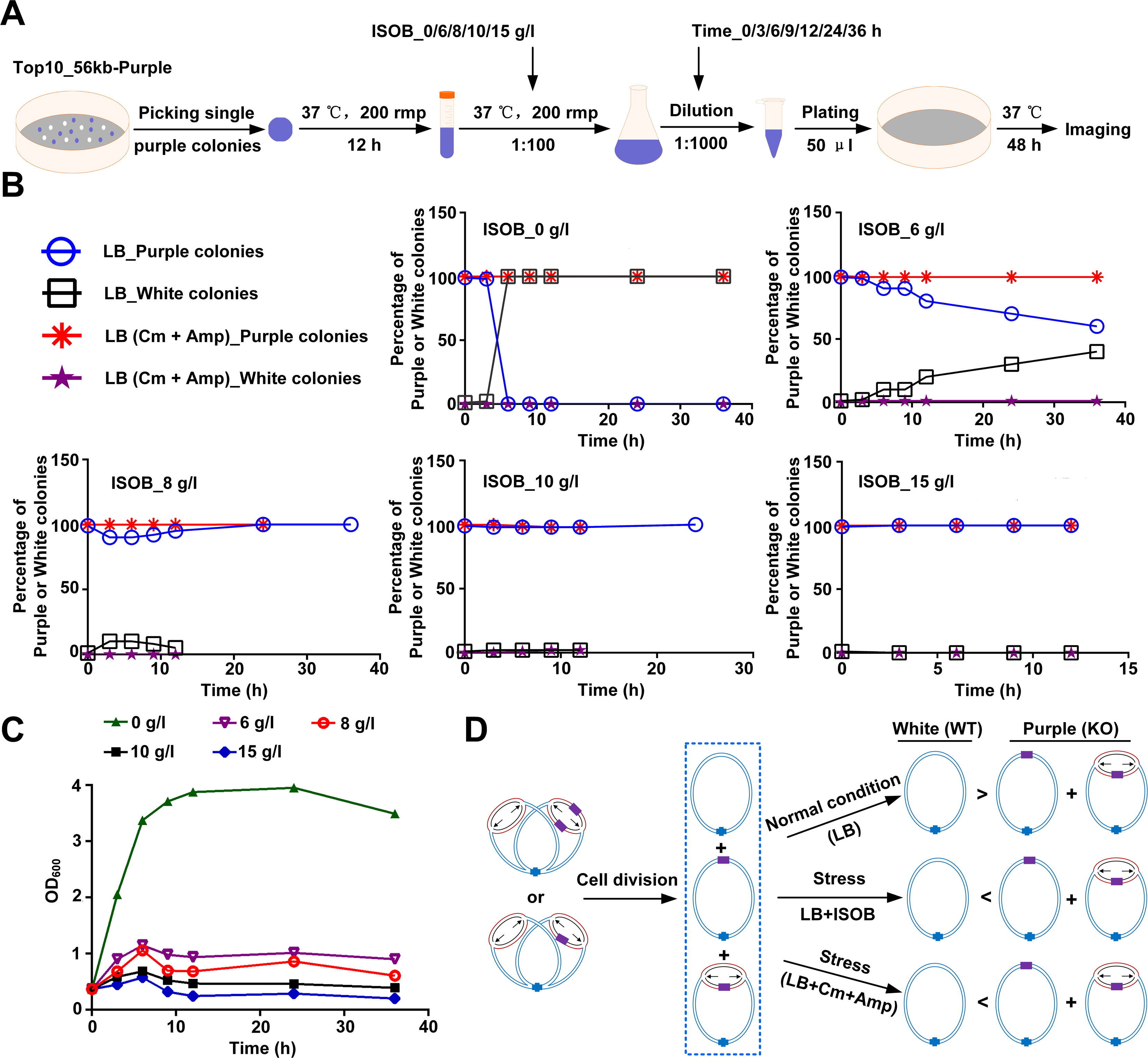
Bacterial local polyploidy or “heterozygote” responded to environmental stress. A An illustration of the Top10_56 kb-Purple inoculated in LB medium containing different concentration of isobutanol to measure response capacity of bacterial or local polyploidy or “heterozygote” to environmental stress. A single colony from LB (Amp + Cm) plate was randomly picked and inoculated in 5 ml LB (Amp + Cm) liquid medium. Cells were cultivated at 37 °C for 12 h. 1% (vol/vol) of the culture was inoculated in 30 ml LB medium in 250-ml baffled shake flasks and grew at 37 °C, followed by the addition of isobutanol with desired concentrations. 1 μl of the culture was taken out and diluted in 1000 μl ddH_2_O. Then, 50 μl of cells were spread on plates with different antibiotics, and cultivated at 37°C. B Top10_56 kb-Purple single colony inoculated in LB medium containing different concentration of isobutanol (closed triangles) could have different phenotypes in the daughter cells. ISOB_6 g/l (See also **Supplementary Figure 27**), 8 g/l (See also **Supplementary Figure 28**), 10 g/l (See also **Supplementary Figure 29**), and 15 g/l (See also **Supplementary Figure 30**). C Comparison of growth with isobutanol stress. Cells were incubated in LB at 37 °C. Time courses for the growth of Top10_56 kb-Purple in the absence of isobutanol (closed triangles) or in the presence of 6 g/l (closed triangles), 8 g/l (closed circles), 10 g/l (closed diamond), and 15 g/l (closed square) isobutanol. D Schematic of bacterial local polyploidy or “heterozygote” responding to isobutanol stress. Described as bacterial local polyploidy or “heterozygote” phenomenon, the same locus at the different copies of chromosomes in the replication fork of a single cell could have different genotypes, such as PPRG and wild-type. In the presence of isobutanol stress, these genes could be allocated into the daughter cells in different combinations, so that the daughter cells have different phenotypes (WT indicates wild-type strain; KO indicates engineered strain).

In order to further study the suitability of the local polyploidy or “heterozygous” purple engineered bacteria Top10_56 kb-Purple to isobutanol and the selectivity of isobutanol to the evolution of the strain, 0-15 g/l isobutanol stress gradient was set at different incubation times to observe the phenotype and growth status of the strain (**Fig. 5b**). First, one single colony of the Top10_50 kb-Purple strain was inoculated into LB (Cm + Amp) liquid medium and cultured at 37 °C as seed culture until OD_600_ reached 0.5. The seed culture developed mostly purple colonies with several white colonies when plated on the LB (no antibiotic) plates, and developed only purple colonies on the LB (Cm + Amp) plates (**Fig. 5b and Supplementary Figure 26**). Then, the seed culture was inoculated into the LB medium containing 0, 6, 8, 10 and 15 g/l isobutanol, respectively, at a rate of 1: 100, and cultured at 37 °C and 200 rpm. At 0, 3, 6, 9, 12 h, sample cultures were plated on LB without antibiotic and LB (Cm + Amp) plates at 37 °C to observe the growth of the colonies (**Fig. 5b and 5c**). Colonies grown on LB media with or without antibiotics showed similar responses toward isobutanol stress. In the absence of isobutanol, the purple cells were quickly replaced by the white ones. In the presence of 6 g/l isobutanol, the purple cells receded gradually while the white ones continued to increase. In the presence of 8 g/l isobutanol, the purple cells were enriched rapidly after an initial decline, and soon dominated the whole population at the expense of white cells. In the presence of 10 g/l or 15 g/l isobutanol, the purple cells remained total dominance but the whole population gradually died out as incubation continued (**Supplementary Figures 26-32**). In this study, the engineered 56 kb deleted strains could survive at least 36 h in 0, 6 and 8 g/l isobutanol, 24 h in 10 g/l of isobutanol, and 12 h in 15 g/l of isobutanol. When plated on LB plates without antibiotic, it was shown that the change of isobutanol concentration could change the ratio of purple (engineered strain) and white colonies (wild-type strain) in the bacterial community (**Supplementary Figure 26**). In the range of 6-8 g/l isobutanol, the number and proportion of purple bacteria were positively correlated with the concentration of isobutanol and the screening time. Isobutanol concentration higher than 10 g/L was toxic even to the engineered cells. We believe that this local polyploid phenomenon of bacteria can act as a protective mechanism to make the bacteria obtain robust properties, avoiding the damage of random mutations and short-term environmental changes to the survival of the whole bacteria population (**Fig. 5d**).

## 4. Discussion and conclusion

### 4.1 Potential impact of multifork on bacterial gene editing

CRISPR/Cas9 technology has been applied to the field of gene editing for just a few years, but its development and utilization surpassed any other recombinant technology such as Cre(39), Flp(40), ZFN(41), TALEN(42). In this study, CRISPR/Cas9 technique was used to localize the PPRG at different positions on the bacterial chromosome, and thus confirming the multifork of bacteria from the single cell level. Our results indicated that the phenomenon of the fork does not only exist in the rapid growth period of the bacterial population (**Supplementary Figure 33**), and the size and duration of the replication fork may also be greater than our traditional belief. Through the multi-round CRISPR/Cas9 editing, the same locus of the different copies of the chromosome in a replication fork in a single-cell genome can have wild-type, PPRG and KRG, and become local polyploidy or “heterozygous” cells. This observed phenomenon suggested that we should pay special attention to the “heterozygous” cell phenomenon caused by multifork when we use CRISPR/Cas9 technology to perform gene editing, especially large fragment gene knockout.

### 4.2 Impact of local polyploidy or “heterozygous” cells in pure homozygote strain screening

In a local polyploidy or “heterozygous” cell, different alleles exist in the same locus and chromosome separation could separate the alleles into two daughter cells. Even in cells proceeding four or even eight rounds of replication simultaneously, theoretically only two or three rounds of chromosome separation could eliminate all local polyploidy or “heterozygous” cells. The astonishing stability of local polyploidy or “heterozygote” phenomenon we observed was most likely due to the HR between different alleles inside the different branches of the multifork chromosome(24). Theoretically, cells with “heterozygote” genotype, although represent a very small proportion of the whole population, could be kept forever in a population through HR. In our experiments, each visible colony in the first two rounds of plating had approximately 10^7^ engineered cells, and were demonstrated to contain some copies of the wild-type alleles in the chromosome by the colony PCR reactions. Since the visible colony took more than 20 rounds of cell generations to form, the local polyploidy or “heterozygous” cells were demonstrated to be able to last even for hundreds of cell generations in a strain population in this study (**Supplementary Figure 25**). Taken together, this continuous culturing experiment showed that it might take hundreds of cell generations to form a pure homozygote strain colony containing absolutely no local polyploidy or “heterozygous” cell, even under the selective pressure conditions. Multi rounds of single-cell plating assays under selective pressure conditions might be needed to obtain a pure population of engineered strains containing absolutely no wild-type allele in the chromosome. In previous genetic engineering experiments, similar “heterozygous” problems, if existed, might be overlooked because the amount of “heterozygous” cells were too small to affect the phenotype of the whole strain population.

### 4.3 Relationship between local polyploidy or “heterozygous” cells and strain degeneration

The qualities of the engineered strains are the key to determine the yield and quality of a fermentation process. The preservation and use of a seed strain, regardless of the method adopted, will change over time, which is generally regarded as the strain degeneration. The traditional explanations of the strain degeneration includes gene recombination, variation of the control of gene expression and enzyme synthesis, infection of certain viruses, interference with cell normal metabolic activity, long and frequent asexual reproduction, gradual accumulation of mutations, etc. In this experiment, we note that the use of CRISPR/Cas9 technology-mediated bacterial gene editing, could (but not necessarily) lead to local polyploidy or “heterozygous” cell phenomenon caused by multifork. The persistence of local polyploidy or “heterozygous” cell, even at a low abundance, in the cell population could result in engineered strain degeneration under certain stressed environmental conditions. To avoid potential risk of strain degeneration, pure homozygote colony without any local polyploidy or “heterozygoos” cell should be stored and used regardless of the gene editing technologies selected for strain engineering.

Taken together, we believed that this bacterial multifork phenomenon provided a pattern of eukaryotic alleles for prokaryotes. This phenomenon could be an intrinsic driving force for the periodic change of the dominant bacterial populations, enabling the bacteria a potential to deal with the unexpected changes in the environment. This phenomenon provided a protective mechanism for the bacteria and the robustness to adapt to the environmental changes, and to avoid random mutation and short environmental changes to damage the bacterial population.

